# Hippocampal replay repairs the cognitive map for flexible planning

**DOI:** 10.64898/2026.10.20.760493

**Authors:** Armin Bazarjani, Caitlin S. Mallory, David J. Foster, Payam Piray

## Abstract

Hippocampal replay is structured: prospective replay typically precedes retrospective replay, each aligns with the animal’s future and past paths, and replay sometimes targets abandoned goals. We propose that replay repairs the cognitive map. A map that supports efficient planning precomputes expected paths; because plans terminate at the goal, the goal is part of the structure it represents. Every goal change therefore leaves the map incorrect and requires two repairs: one for the goal adopted, one for the goal left behind. The larger is expressed first, and the path each takes determines what it depicts, accounting for replay’s ordering, direction, and paradoxical content. The model also generates the predictions that this ordering should invert when past and future paths converge. Recordings from rats foraging in an open arena confirm this: replay order is not as fixed as previously believed, but varies with trajectory geometry and the proximity of the abandoned goal.

## Introduction

Hippocampal replay is highly structured. During pauses in behavior, place cells reactivate in temporally compressed sequences that trace trajectories through the environment, typically extending away from the animal’s current location [1–5]. Within a single stopping period, prospective replay reliably precedes retrospective replay across mazes and sessions and even in animals with minimal experience of the environment [6]; prospective replay aligns with the path the animal is about to take and retrospective replay with the path it has just taken [2, 7, 8]; and the balance between the two directions shifts with the demands of the task [9]. The content of replay is equally orderly, though in ways that are harder to interpret. Replay can depict the precise path to the animal’s current goal moments before the animal takes it [5, 10], but it can also depict the goal the animal has just abandoned, or the arm it will not choose, in a manner anti-correlated with behavior [11–14]. These regularities are the most direct evidence available about what replay is for. However, taken together, they have been difficult to attribute to a single computational process, since replay sometimes appears to serve the animal’s next decision and sometimes to work against it.

A cognitive map that is to be used for planning must be kept current, a requirement which has so far received little attention. Planning at decision time is fast only if the map has already been compiled for the task the animal faces. In episodic decision problems, the goal enters that compilation as a terminal state, i.e. a state at which the episode ends and beyond which the plan does not extend. Terminality is therefore part of the transition structure the map represents, not an annotation on top of it. Importantly, each time the animal’s goal shifts, the reassignment of terminal states alters that structure. A map compiled for the previous goal is stale for the next one, and of everything that can change in an animal’s world, its goals change most often. Much work has addressed how cognitive maps are constructed in the first place [15–17]; much less has asked what keeps an existing map usable as goals change [18–20]. The question is a natural one to put to replay, which occurs precisely in the intervals between goal-directed episodes, when the map must be brought up to date before it is next consulted.

Here, we propose that replay repairs the hippocampal cognitive map. We formalize the map as the default representation (DR) [18], a predictive map [21] that supports planning in closed form, without search at decision time. Crucially, when the transition structure changes, the DR can be repaired exactly by a low-rank update so that the map is updated in place rather than relearned [18, 19]. We hypothesize that this repair is the computation carried out by replay. The hypothesis is specific enough to determine which updates a goal switch requires, what trajectory each update traces, and which of them should be expressed first. For a goal switch, the repair decomposes into exactly two updates, one that makes the upcoming goal terminal and one that makes the previous goal nonterminal. We identify these with prospective and retrospective replay respectively. Importantly, each repair is applied state by state, with successive states drawn from the update matrix itself, so that it unfolds as a trajectory through the state space. That trajectory is what is expressed in replay. In this framework, the priority of each update is given by its magnitude, which, because the update is rank one, is the absolute value of its single nonzero eigenvalue.

This single computational mechanism, with no tuning between tasks, accounts for findings that have so far required separate explanations. Because imposing terminality on a state perturbs the map more than removing it, the prospective update is the larger of the two and is selected first, reproducing the prospective-before-retrospective ordering reported by Mallory et al. [6]. Moreover, since this ordering follows from the structure of the updates (rather than from learning), the model also predicts it in animals with little experience of the environment, consistent with empirical data. Because each update propagates along the paths connecting the map to the goal it concerns, prospective replay aligns with the future path and retrospective replay with the past path [8]. Paradoxical replay follows as well: an obsolete goal must be stripped of its terminality before the map can support planning to any other location, so replay is directed at the goal the animal has abandoned, or at the arm it is not choosing [12, 13]. The same reasoning explains why replay appears to lead behavior in one T-maze design and to lag it in another [10, 13], since the two tasks require different repairs.

The account also makes a novel prediction, that retrospective replay should vary across trials in a specific way that has not previously been tested. The reason is that the approximation behind the ordering does not hold equally on every trial, since the two repairs interact when their paths overlap. Specifically, the eigenvalue of each repair sets which comes first, and it is also what changes when the approximation fails. When the animal’s past and future paths converge, the two repairs run along much the same stretch of the map, so the prospective repair is computed on a map that still treats the previous goal as an endpoint, and its eigenvalue is reduced; the closer the two goals lie, the greater the reduction. When the paths diverge, the repairs act on separate parts of the map, the approximation holds, and the prospective repair stays the larger however near the previous goal happens to be. Retrospective replay should therefore increase with proximity to the previous goal when the paths converge, and should be unrelated to proximity when they diverge. We tested these predictions in an existing dataset of hippocampal ensemble recordings from rats foraging in an open arena, where reward alternated between a fixed home well and an unpredictable away well, and where in most sessions the animals had to navigate around transparent barriers [6]. Both predictions were confirmed. On converging-path trials, the rate of retrospective replay (normalized by the duration of each stopping period) increased steadily with proximity to the previous reward well; on diverging-path trials, it was unrelated to distance. This rules out the simpler possibility that replay merely favors locations near the animal.

On this view, replay maintains the representation that planning relies on, and its goal-directed and anti-goal-directed forms reflect the same computation applied to whichever part of the map is currently wrong.

## Results

### Repairing a predictive map

#### The predictive map

We represent the cognitive map as the default representation (DR) [18], one of a family of predictive maps [22, 23] proposed for the hippocampus [21]. The map is an *N* × *N* matrix **D**, where *N* is the number of states, and each entry encodes the expected aggregate cost of all future visits to *s*^′^ from *s*:

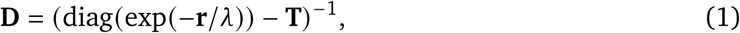

where **r** is the reward vector across all states, with non-zero rewards at terminal states, and **T** is the *N* × *N* transition matrix, whose entry *T_i j_* gives the probability of moving from state *i* to state *j* in one step. Because the optimal value function follows from **D** in closed form (Methods), planning with this map requires no search at decision time, which is what makes it a candidate for a map an animal could exploit at a decision point. Moreover, since equation (1) inverts a matrix built from **T** and **r**, any change to either invalidates the map compiled from them. A goal is what reinforcement learning theory calls a terminal state: a state the agent does not leave, whose row of **T** is replaced by a self-transition (one on the diagonal, zero elsewhere). Making a state terminal therefore changes **T**, and so does making a terminal state nonterminal again. A goal switch requires both, and so invalidates the map.

#### Repairing the map

A map defined by equation (1) need not be recomputed when the environment changes. Local changes to the transition graph can instead be absorbed by an in-place correction [18, 19]:

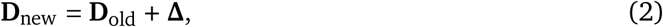

where **Δ** is a low-rank change matrix whose rank equals the number of states with modified transitions, and which is cheap to compute (Methods). The correction is exact: the repaired map is identical to the one obtained by recomputing equation (1) from scratch. A goal switch requires two such corrections, one making the new goal terminal and another making the previous goal nonterminal. Each repair is computed from the map in place before either has been applied, thus each treats the change it concerns as the only change to the map. This is accurate when the two corrections act on well-separated parts of the map, and a first-order approximation when they do not [19].

#### Repair as a trajectory

Equation (2) is written as a single correction to the whole matrix, which is biologically and psychologically implausible, since it requires simultaneous access to the relation between every pair of states, including states the animal is neither occupying nor considering. But the correction is defined separately for each state, so each row of the map can be corrected on its own,

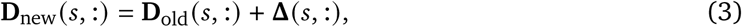

for any state *s*, leaving the map corrected wherever the update has been applied and unchanged elsewhere. Therefore, a system that can visit states only one at a time is still able to carry out the repair, as long as it visits the right states.

This raises the question of which states should be visited, and in what order. Importantly, the update itself already contains the information needed, and so provides an answer. Specifically, the row **Δ**(*s*, :) records how much the relation between *s* and every other state has been disrupted by the change, and these entries are largest for the states connected to *s* through the region the change actually affects. Therefore, a repair that moves each time to the state with the largest entry is following the change outward from where it originated, correcting each state’s relations before it reaches the states whose relations depend on them. Accordingly, we take the next state to be the one with the largest entry in **Δ**(*s*, :). The repair then traverses the state space along a path dictated by the structure of the change, and it is this path that we identify with a replayed trajectory. Two properties of a repair follow, and both are used throughout what follows: the path it takes depends on which state’s terminality is being changed, and its magnitude, which we take as the single nonzero eigenvalue *λ*_0_ of **Δ**, depends on how much of the map the change disturbs.

#### Repaired maps support planning

Before using this scheme to model replay, we checked that a map repaired over a subset of states remains usable. We built the DR for a maze in which the only rewarding terminal state was in one corner, then moved the goal, which requires making the new location terminal and the previous one nonterminal, and applied the two corrections state by state as in equation (3). Planning with the repaired map recovered the shortest path to the new goal, and did so as well as planning with a map reconstructed in full from equation (1) (Supplementary Figure 1). The same holds when the goal moves repeatedly, as it does in tasks where an animal must reach an intermediate location before a final one.

### Prospective replay precedes retrospective replay because its repair is larger

Both repairs called for by a goal switch are needed, so the question is which is carried out first. They are not equal in size. Because a single change in terminal status modifies the transitions of one state, each repair is a rank-one matrix, and its magnitude is given by the absolute value of its single nonzero eigenvalue, |*λ*_0_|, which measures how much the change disturbs the map overall. Imposing terminality on a state disturbs the map more than removing it from another, and across our simulations the prospective repair is consistently the larger of the two. Probabilistically selecting between the two replays over their respective magnitudes therefore yields the prospective repair first (Figure 2**c**,**d**). Note that this asymmetry is a property of the two repairs and not of the animal’s history, so the same ordering is expected whatever the animal’s experience of the environment.

**Fig. 1.**
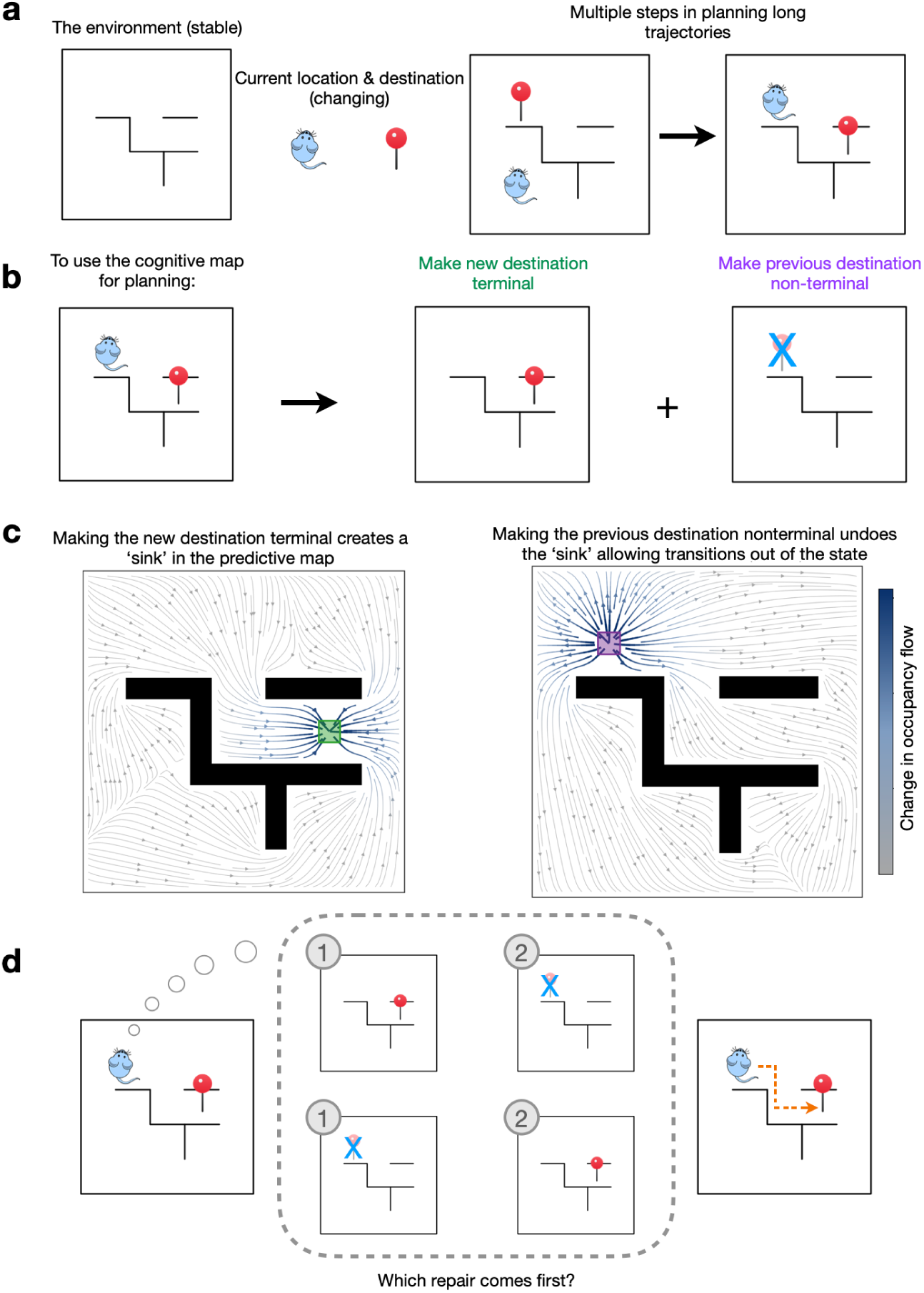
The map-repair model of replay. **(a)** High-level overview. The environment provides a stable representation while the current location of the animal and the desired destination are dynamic and always changing as the animal moves throughout the environment taking long sequences of actions. **(b)** The map needs to be repaired in two steps to be used for planning. One repair makes the new destination terminal, that is, a state at which the animal’s plan ends; the second makes the previous destination nonterminal again, so that plans can pass through it. **(c)** Visualization of what these repairs do to the map. Making a state terminal turns it into a *sink*, making trajectories flow into it and not out. Making a state nonterminal undoes this operation and now allows trajectories to flow out of the state. **(d)** The two repairs are applied sequentially with one repair being applied after the other. They can be applied in either order with the larger repair being selected first, the size of the repair depends on how much of the map it changes.

**Fig. 2.**
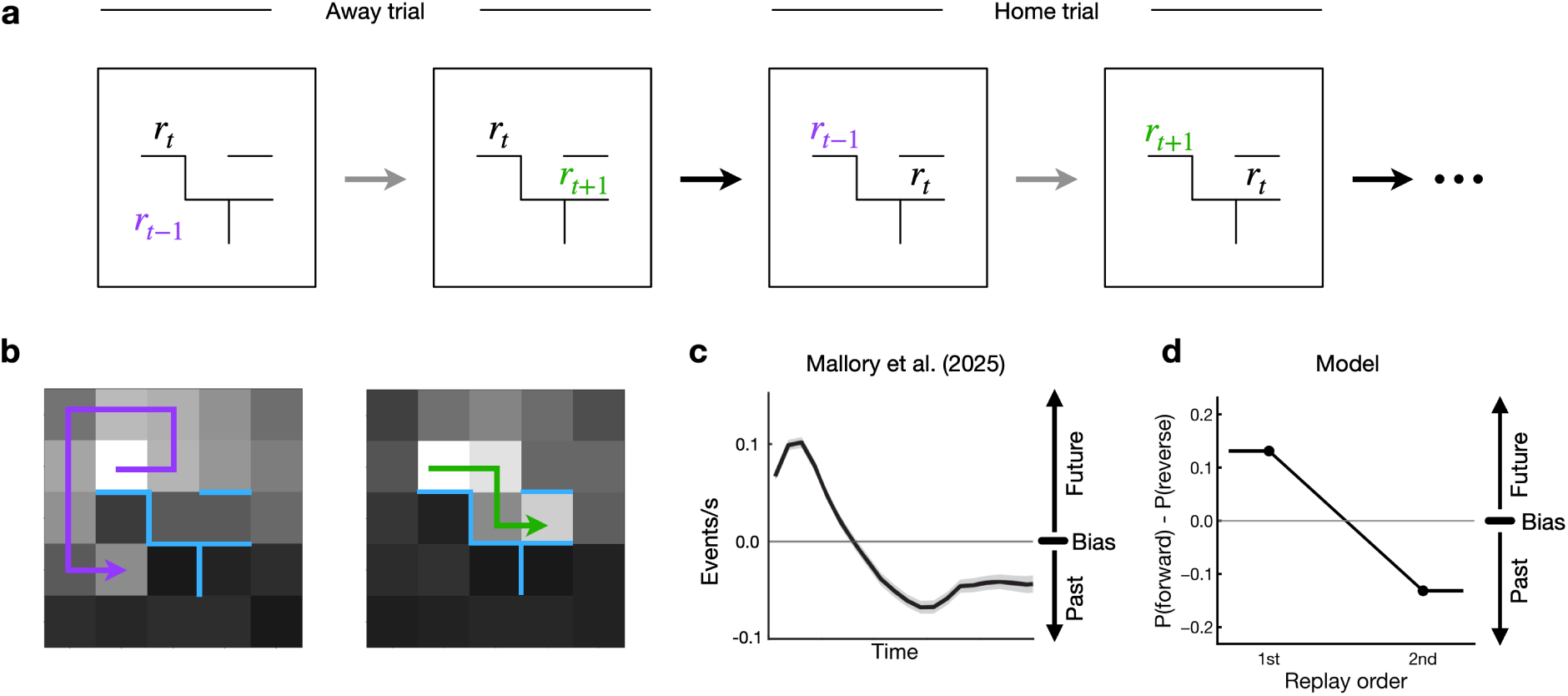
Prospective replay is prioritized over retrospective replay. **(a)** Open-field maze on which we tested the model. *r_t_*_−1_ denotes the location the rat traveled from to reach the current state, *r_t_* is the rat’s current state, and *r_t_*_+1_ is the next goal state the rat will travel to. The purple line shows the rat’s previous path from *r_t_*_−1_ → *r_t_* and the green line shows the future path from *r_t_* → *r_t_*_+1_. **(b)** Visualization of prospective and retrospective replay: the update required to make the future state terminal is more aligned with the agent’s future trajectory (prospective replay), while the update required to make the current goal state nonterminal is more aligned with the agent’s previous trajectory (retrospective replay). **(c,d)** Timecourse of replay: **(c)** Across conditions, replay is initially biased towards the future (prospective) and then the past (retrospective). **(d)** The model reproduces this preference for prospective first, then retrospective. Bias is the probability of carrying out the prospective first minus that of the retrospective repair. The model specifies the order of the two replays rather than their timing. Data in **(c)** adapted from Mallory et al. [6].

This is what is observed. Within a stopping period, prospective replay is preferentially expressed before retrospective replay, on a linear track and in an open field alike, shortly after the animal stops and almost always preceding retrospective replays of the path just taken [6]. The ordering holds across subjects, sessions, and stopping periods containing multiple events, and it is present in animals with limited experience of the environment, which suggests a mechanism that is intrinsically generated rather than learned. On our account it is intrinsically generated in a specific sense: it reflects the relative size of two corrections that any goal switch requires.

### Prospective and retrospective replay follow the paths their repairs take

The two repairs also act on different parts of the map. The repair that makes the upcoming goal terminal acts on the states from which that goal is reachable, so it propagates along the paths leading into it. The repair that makes the previous goal nonterminal acts on the states around the goal being abandoned, so it propagates along the paths leading out of it. Whichever repair is being carried out, the trajectory it traverses is therefore determined by the goal it concerns: the prospective repair should follow the animal’s future path and the retrospective repair its past path. This is a consequence of a single mechanism, not of two mechanisms with separate functions.

The prediction is borne out at decision points, where past and future paths are distinct and behaviorally defined. Shin et al. [8] recorded replay in rats performing continuous spatial alternation on a W-shaped maze, in which inbound trajectories run from a side well to the center and outbound trajectories from the center to the opposite side well. Replay direction tracked behavior rather than maze location: prospective replay represented outbound paths at the center well and inbound paths at the side wells, while retrospective replay represented inbound paths at the center well and outbound paths at the side wells. Simulating the same task reproduces this pattern (Figure 3**c**). At the center well the retrospective repair traverses the past inbound trajectory and the prospective repair the future outbound trajectory; at the side wells the relationship inverts, with the retrospective repair traversing the past outbound path and the prospective repair the future inbound path.

**Fig. 3.**
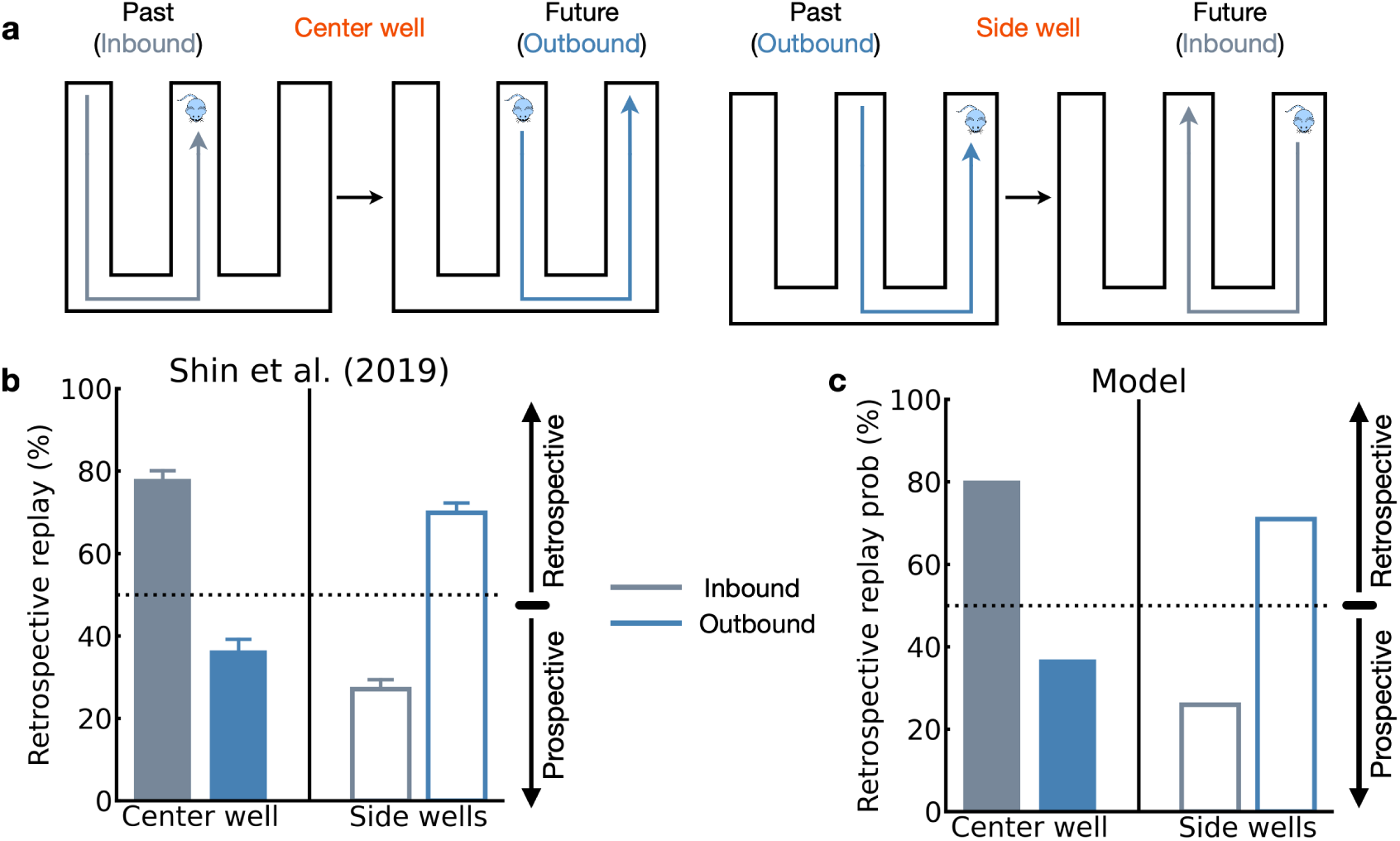
Prospective replay of future paths, retrospective replay of past paths. **(a)** Task design. Rats were trained to run through a W-maze, receiving reward for successful completion of two types of runs, center-side-center and side-center-side. **(b)** Data replotted from Shin et al. [8] showing the percentage of retrospective replays associated with the inbound and outbound paths, and the percentage of prospective replays associated with the outbound and inbound paths, at the center and side wells respectively. **(c)** Probability of the retrospective repair replaying the inbound path (light blue) and outbound path (dark blue), and probability of the prospective repair replaying the outbound path and inbound path, at the center and side wells respectively. Data in **(b)** adapted from Shin et al. [8].

### Replay looks paradoxical because it is directed at abandoned goals, where the map is wrong

Replay often reactivates paths the animal is not about to take, and sometimes paths it has deliberately stopped taking [12, 13]. This has been called “paradoxical” [14], since it runs against the expectation that replay is directed at the animal’s current goal. On our account it is the expected consequence of a goal that has become obsolete. A state the map still treats as terminal absorbs any plan that passes through it, so a goal the animal has abandoned must be made nonterminal before the map can support planning to anything else. Replay should therefore be directed at abandoned goals, not because the animal intends to return to them, but because that is where the map is wrong. Two well-known results follow from this.

In the first [12], rats foraged in an eight-arm maze in which one arm dispensed reward until, after a fixed number of visits, it stopped and another arm became rewarding. Each block therefore had a search phase, in which the rats had to find the new arm, and a repeat phase, in which they returned to it (Figure 4**a**). Throughout both phases, replay preferentially represented the path to the previous, now unrewarding, arm (Figure 4**b**). Under our account, that arm’s terminal status is obsolete as soon as the rat learns it no longer pays, and stripping it is what the search phase requires of the map: the rat has entered a phase in which the remaining seven arms must be explored, and the map cannot support planning to any of them while it still treats the old arm as an endpoint. Replay should be directed there, and in the model it is (Figure 4**c**).

**Fig. 4.**
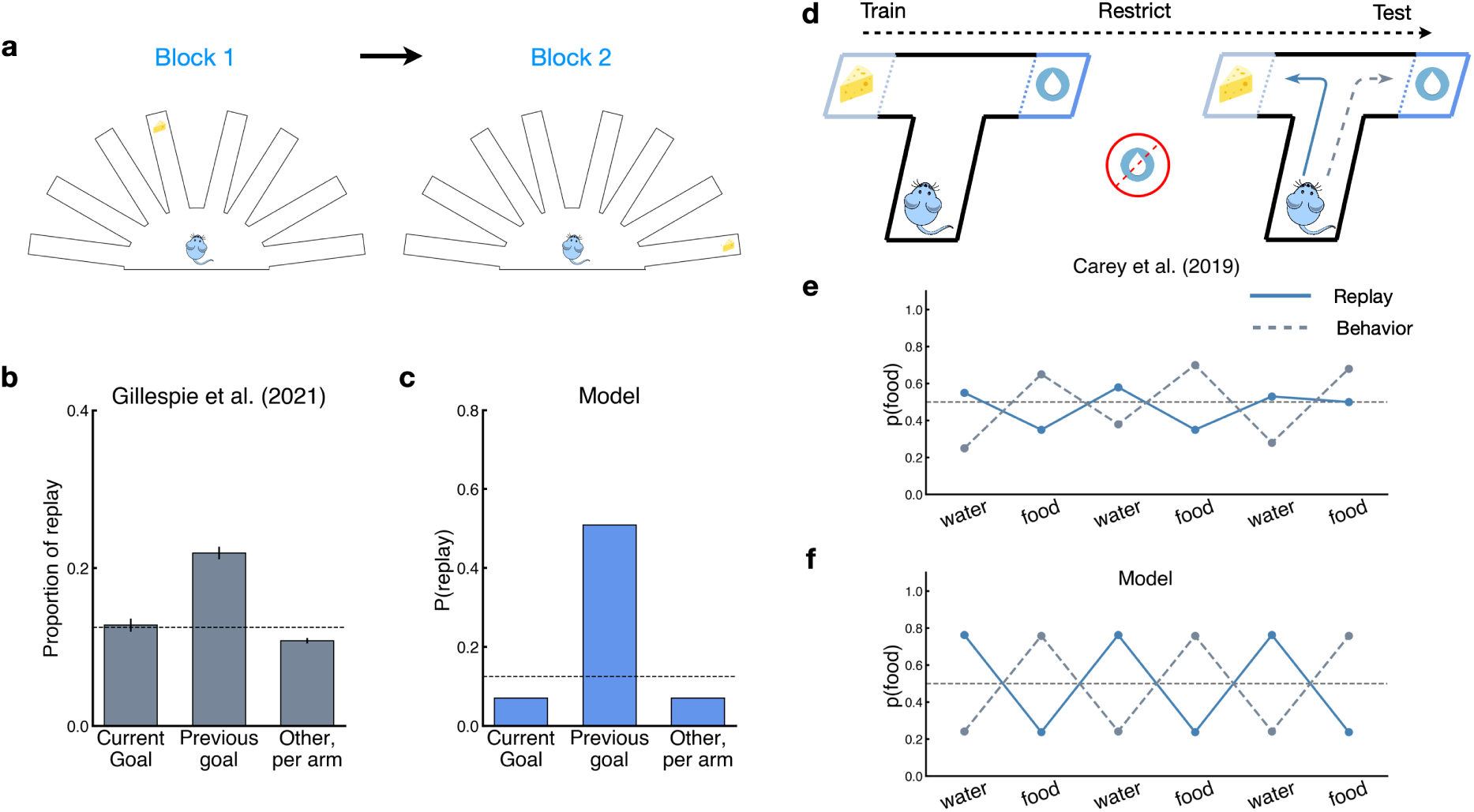
Paradoxical replay. **(a)** Task design from Gillespie et al. [12]. Rats learned that when a goal appeared in an arm, that arm would be continuously rewarding for a period of time. Once the arm became unrewarding, the rats had to search for the new arm containing reward. **(b)** Data replotted from Gillespie et al. [12] showing the proportion of replay towards the current goal, the previous goal, and every other arm. Replay preferentially goes towards the previous goal arm. **(c)** Probability of replay going towards the different arms, computed from the model. **(d)** Task design from Carey et al. [13]. Rats are placed in a T-maze and learn that each arm contains a distinct reward (either food or water). They are then restricted access to one of the two resources and placed back into the maze. **(e)** Data replotted from Carey et al. [13] showing the proportion of replay events going towards the food arm (blue) and the proportion of trials on which the rat went to the food arm (gray). Replay is anticorrelated with behavior: it preferentially goes towards the restricted resource, opposite to the rat’s own behavior. **(f)** Probability of replay going towards the food arm (blue) and probability of the agent going towards the food arm (gray), computed from the model. Data in **(b)** adapted from Gillespie et al. [12]. Data in **(e)** adapted from Carey et al. [13].

In the second [13], both arms of a T-maze contained a distinct reward, food in one and water in the other, and rats foraged freely until they had learned the association. The experimenters then restricted access to food or water, inducing hunger or thirst, and returned the rat to the maze (Figure 4**d**). Behavior followed the restricted resource, but replay ran the other way, toward the arm the animal was not choosing, so that replay was anti-correlated with behavior (Figure 4**e**). Here both arms hold their own standing reward and the rat is aware of both, so the map represents both as terminal. That is the correct long-run representation of this task, since both arms remain available throughout and each reward is unchanging. But a map with two terminal states cannot be used to plan to either one. Planning to the preferred arm requires making the other nonterminal, and the repair that does so is directed at the arm the animal is not choosing (Figure 4**f**).

The same reasoning resolves a discrepancy between two superficially similar T-maze experiments. Replay appears to lead behavior in one [10] and to lag it in the other [13], and the difference lies in what each task makes terminal. In Ólafsdóttir et al. [10] the arms were inaccessible before test and so were not terminal; when the cued arm was revealed, the map required that arm to be made terminal, which is a prospective repair and looks goal-directed. In Carey et al. [13] both arms were already terminal, so the required repair was the removal of terminal status from one of them, which is retrospective and looks paradoxical. Simulating both tasks in the same environment under these two schemas reproduces both patterns (Supplementary Figure 3).

### Retrospective replay is prioritized when its repair magnitude becomes the larger of the two

The results so far show the prospective repair to be the larger of the two, which is what places prospective replay first. This appears to commit the model to that ordering unconditionally. However, that rests on an approximation. Specifically, both repairs are computed from the same map, the one in place before either has been applied, so each is evaluated as though the other had not happened. This approximation is accurate as long as the two repairs do not touch the same states. Moreover, since each repair is applied stepwise along a path, it updates only the states on that specific path, so the two interact mainly when their paths overlap. As long as the two paths do not overlap, the repairs act on largely separate parts of the map, which means the approximation is accurate, and the prospective repair remains the larger. When they do overlap, by contrast, the prospective repair is computed on a map that still treats the previous goal as an endpoint, a map the retrospective repair, when it is carried out, will correct along the very states the prospective repair will traverse. Thus, while the prospective repair’s magnitude is reduced, the retrospective repair, which concerns the state the animal already occupies, remains relatively unaffected. Furthermore, the closer the two goals lie, the larger this reduction will be, since more of the prospective repair’s path then passes through territory the retrospective repair also covers. Notably, the paths overlap, and the goals lie close, exactly when the animal is turning back toward where it came from. When the reduction is large enough, therefore, the prospective repair falls below the retrospective one and the ordering inverts.

Retrospective replay should therefore increase as the previous well lies closer to the animal when the past and future paths overlap, and show little or no such increase when they diverge. Simulating well configurations ranked by their optimal distance from the previous well gives this pattern: on converging-path trials the probability of selecting the retrospective repair rises steeply as the previous well approaches (Figure 5**e**); on diverging-path trials it remains near one half at every distance, with no comparable rise (Figure 5**f**). These simulations use the same model, with the same parameter values, as those reported above.

**Fig. 5.**
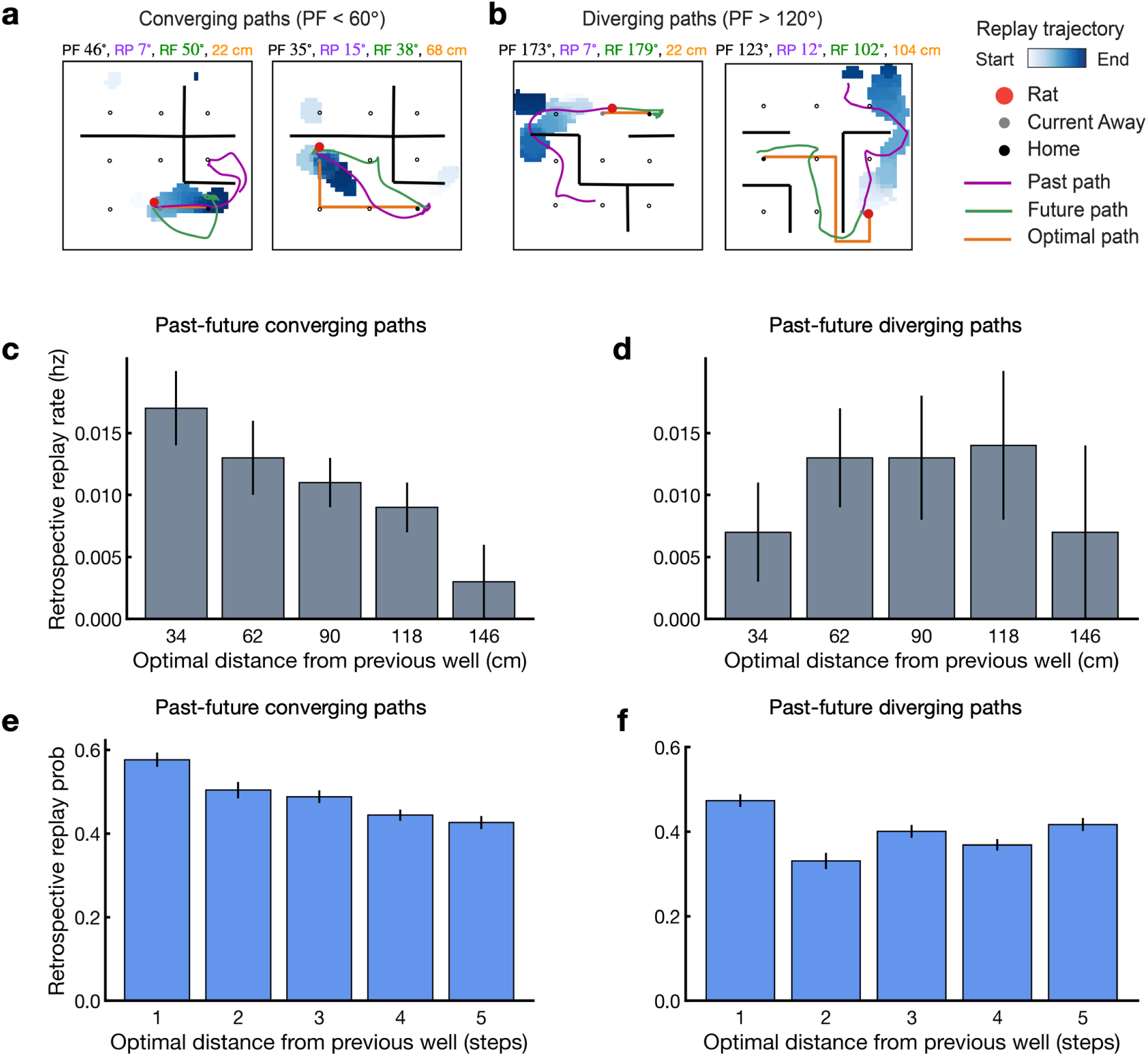
Map repair predicts when retrospective replay is prioritized. **(a, b)** Examples of retrospective replay events, from our analysis of the recordings of Mallory et al. [6], detected on trials in which the past and future paths converged (**a**) or diverged (**b**). For each path configuration, examples are shown for trials in which the future well was relatively close to (left) or far from (right) the rat. The optimal path distance from the rat to the future well is indicated in orange above each example. PF, past-future angular displacement; RP, replay-past angular displacement; RF, replay-future angular displacement. **(c, d)** New analysis of the recordings of Mallory et al. [6]. Rate of retrospective replay as a function of the optimal path distance from the previous reward well, on away trials with converging (< 60^◦^; **c**) and diverging (> 120^◦^; **d**) past and future paths. Retrospective replay rate declines with distance when the paths overlap, and shows no systematic relationship when they diverge. **(e, f)** Simulations of the same task. Probability of selecting the retrospective repair over its prospective counterpart, for well configurations sorted by optimal distance from the previous well, on converging (**e**) and diverging (**f**) trials. Error bars denote s.e.m. across trials.

Is this true of hippocampal replay? In recordings from rats navigating an open arena for reward delivered at a consistent home well on alternating trials and at an unpredictable away well otherwise [6], the rate of retrospective replay rises as the previous well lies closer to the animal on trials where the past and future paths overlap (Figure 5**c**), and shows no comparable rise on trials where they diverge (Figure 5**d**). Analyses were restricted to away trials, on which the rewarded well was drawn at random, so that the geometry relating the previous, current and next wells varied from trial to trial (Methods).

To confirm this statistically, we modeled the number of retrospective replay events emitted during each stopping period as a Poisson variable, using the log of the duration of the stopping period as an offset so that the coefficients are interpretable as effects on replay rate rather than count. We first asked whether the influence of distance on retrospective replay depends on the geometry of the past and future paths. It does: retrospective replay rate declined with the optimal path distance between the previous well and the current well (*β* = −0.019 ± 0.005, *p* = 3.8 × 10^−5^), and this decline was significantly modulated by the past–future path angle (interaction *β* = 0.0083 ± 0.0031, *p* = 0.0080; *n* = 1908 stopping periods), with the negative effect of distance weakening as the past and future paths diverged and vanishing at an angle of ∼129^◦^.

Fitting the two regimes separately confirmed this. On converging-path trials (past–future angle < 60^◦^), retrospective replay rate fell steeply with distance from the previous well (*β* = −0.0109 ± 0.0030, *p* = 2.9 × 10^−4^; *n* = 925), corresponding to a ∼10% reduction in retrospective replay rate for every additional 10 cm and a ∼70% reduction across the range of distances sampled (34–146 cm). On diverging-path trials (angle > 120^◦^), the same model returned an estimate close to zero (*β* = −0.0004 ± 0.0051, *p* = 0.93; *n* = 339; 95% CI on the rate ratio per 10 cm: [0.90, 1.10]), so that the data admit at most a small effect of distance in this condition, as the model predicts. The same pattern held with non-parametric tests that make no assumption about the form of the relationship: splitting converging-path trials at the terciles of the distance distribution, retrospective replay rates were higher when the previous well was close than when it was far (rank-sum *z* = 2.81, *p* = 0.0049), and retrospective replay rate correlated negatively with distance across the full range (*ρ* = −0.088, *p* = 0.0078). Neither effect was present for diverging paths (rank-sum *z* = −0.04, *p* = 0.97; *ρ* = 0.010, *p* = 0.86).

The difference between the two conditions is what distinguishes this account from a simpler one on which replay favors whatever lies near the animal, since such a bias would produce a distance effect of the same size whatever the path geometry. What sets the size of the retrospective repair is how close the abandoned goal lies to the animal, and what sets whether that repair is large enough to be carried out first is how close the two goals lie to each other.

## Discussion

We have argued that the regularities of hippocampal replay, its ordering within a stopping period, its alignment with past and future paths, and its frequent direction toward abandoned goals, follow from a single requirement. Specifically, a cognitive map that supports planning must be compiled against the current goal, and since the goal changes constantly, the map is stale on nearly every trial. Repairing it requires two corrections, one that makes the upcoming goal terminal and one that makes the previous goal nonterminal. Importantly, every aspect of replay’s structure that we have considered follows from the relative size and location of these two corrections. The larger correction is carried out first, which is prospective replay. Moreover, each correction follows the path through the states it changes, which is why prospective replay traces the future path and retrospective replay the past. Furthermore, the correction that removes an abandoned goal is directed at that goal regardless of whether the animal intends to return, which is why replay sometimes appears paradoxical. Crucially, the same requirement predicted the conditions under which this ordering should reverse, and analysis of hippocampal recordings confirmed this prediction.

None of the existing formal accounts of replay treat the map as something that can go stale. Specifically, in each of them the map is the substrate on which replay acts, where replay is a learning process that runs over an existing map, and what the model supplies is a rule for deciding which experiences are worth learning from. The most influential such rule is that of Mattar and Daw [24], in which each replay performs a value backup and trajectories are prioritized by the expected improvement in future reward. However, since this ties replay to the improvement of impending choices, replay that lags behind or opposes behavior is anomalous under it. Moreover, the ordering it implies within a stopping period runs counter to the one observed. This is because the gain from a backup is largest at reward receipt, which favors retrospective replay first, while the need for one is largest before departure, which favors prospective replay last. More recently, Sagiv et al. [25] generalized the backup over a distribution of possible future goals, so that value-like and map-like replay emerge as extremes of a spectrum depending on the animal’s beliefs about goal statistics. Importantly, this recovers replay toward abandoned goals in the tasks of Gillespie et al. [12] and Carey et al. [13]. However, the recovery is carried by a separately learned, slowly updating estimate of which goals tend to recur rather than by the map, and it also requires a different assumption about the animal’s beliefs in each task. In contrast, under the repair account, the same replay content follows from the map alone, by the same mechanism in every task. That said, the goal-marginalized account does reproduce the gradual rise of replay toward a newly found arm over a block [12], a dynamic that repair in its present form does not produce.

The nearest neighbor of the present account is the proposal that replay constructs the map, binding reusable structural elements such as walls, objects and rewards into memories at remote locations [26], part of a broader line of work on replay as constructive computation [27, 28]. Importantly, both that proposal and ours treat replay as an operation on the map rather than as learning over it. However, they differ in what the operation is for. Specifically, construction concerns building the map when structure is discovered, and therefore the replayed trajectory is explicitly irrelevant there, since a random trajectory builds the map as well as a structured one. In contrast, repair concerns keeping the map usable when the task changes, which in most experiments happens on every trial, and here the trajectory is the path of the repair, which is why the account speaks to ordering and direction. Moreover, the two are complementary at the level of the map itself. Specifically, a low-rank correction is a compositional primitive [19], and the goal switch is the instance of it that behavior demands most often. Thus, the constructive account describes how the map is assembled, whereas the present one describes the operation that keeps it current, and furthermore supplies the machinery for goal switching that the constructive account has been noted to lack [25].

The account requires that the system know which state to make terminal and which to make nonterminal, but it does not say where that knowledge comes from. One plausible source is the prefrontal cortex (PFC). Indeed, medial PFC has long been implicated in maintaining task schemas [29, 30] and in using contextual information to select among competing hippocampal memories [31–34]. Moreover, its damage impairs switching between perceptual dimensions [35], between place and response strategies [36], and between odor and place memories [37]. Thus, medial PFC might specify the current goal in an abstract fashion, such as a reward type or a contextual feature, but not as a location, and the hippocampus converts this into a state of the map, plausibly via reward-vector cells [38] or context-sensitive place cells. In the task of Carey et al. [13], for example, medial PFC registers the shift from food to water, and the hippocampus determines which well thereby loses terminal status and which acquires it. On this view, the coordination between medial PFC and hippocampus observed at decision points [8, 39] is the moment at which a goal switch is communicated, and it likely occurs at the point where it is needed, before the map is next used. The same view fits the finding that hippocampal sequences follow the animal’s current action plan rather than external cues [40], since on our account the sequence is the repair that plan requires. Finally, we note that the repair itself need not be computed in closed form. Specifically, by separating the map into a stable feature basis and an operator that encodes the current environment, one can obtain a temporal-difference rule that recovers the operator from local experience and converges to the exact correction [41]. The same separation also accounts for the stability of entorhinal grid fields across environments alongside their local remapping under local change [19, 41], which in turn suggests where the basis and the correction might respectively be represented.

We would also like to highlight a computational point about why replay is a suitable neural mechanism for implementing repair. Replay has generally been treated in normative models as an opportunity for computation that could in principle happen at any time [24, 42]. However, the situation is fundamentally different for the repair account. Specifically, the states whose rows are corrected are states the animal does not occupy, and replay is known to reach such states [43]. Online experience cannot substitute for this, because a correction that relied on sampling would require the animal to visit each affected location before the map around it became usable again. The repair account does, however, operate under a deadline, because the map has to be correct with respect to its most important element (i.e., the goal) before it is used next time, and that happens the moment the animal departs [23, 44]. Therefore, the interval between arriving at one goal and leaving for the next is not only a convenient time to compute, it is also the window in which the repair has to be done. In this sense, the offline character of replay is not incidental to the computation but a fundamental property of it.

The proposed model has two limitations. First, since each repair is exact and applied once, the model does not generate replay dynamics over time, and therefore cannot address how replay to a given location rises or falls across a block, the pattern noted above that a goal-marginalized backup model reproduces through a forgetting term and a slow learning rate. Nevertheless, the repair account does suggest what the dynamics should look like. Once a new goal arm is found, the repair that makes it terminal is required, so prospective replay toward it should increase. Producing this as a time course would require a variant of the model in which repairs are applied incrementally using a learning rate. Although straightforward, such an extension remains to be built. Second, in some tasks the terminal states follow from the structure of the task, for example when both arms of a T-maze are rewarded. In these cases, we supply that structure in our simulations, so how the animal identifies which state should be terminal remains open. The prefrontal proposal above may be relevant here, and a model that learned task schemas from experience and used them to select the repair would be a natural extension.

Nothing in the mechanism is specific to goals. Indeed, a repair is a low-rank correction for any change in the transition structure, and barriers, doors and objects are such changes [18, 19]. We have concentrated on goals because that is what changes most often in the tasks used to study replay, and because it is the change the replay literature has most directly measured. What we have tested is therefore awake replay in tasks with defined goals. Accordingly, sleep replay, and replay in tasks with no goal change, lie outside that scope. Nevertheless, the proposed framework makes a clear prediction about barriers. Specifically, a barrier should elicit replay with the same organization as goal changes. The trajectory, in particular, should propagate from the altered states, with its magnitude should depend on how much of the map the change disturbs. Moreover, inserting a barrier should differ from removing one in the way that making a state terminal differs from making it nonterminal. Notably, replay does reroute around newly introduced barriers [45]. On our account, its direction, ordering and rate should largely follow the structure of the correction. Another case is human replay. Replay-like sequences reorganize experience according to learned task structure [46] and are expressed during rest in a way that predicts subsequent planning [47, 48], and the account predicts that in tasks where the goal changes, their content should track the repair the new goal requires.

Two further predictions concern goal-related replay itself. First, the size of a repair should also set how much replay it receives, which means that directions that require larger corrections should be replayed more often. Importantly, in principle, a version of the model with incremental learning, which would produce multiple replays per repair, could be tested against replay counts and against the replay of remote locations observed during learning [11]. Second, retrospective replay should precede prospective replay in tasks where the required repair is retrospective, as in Carey et al. [13]. Accordingly, the two T-maze studies compared above [10, 13] should differ in exactly this respect.

Replay has been studied for two decades as a window onto what the hippocampus computes while the animal is not moving. Ask instead what the cognitive map has to be if an animal is to plan with it, and much of the organization of replay follows from the answer. Critically, goals belong to the structure of the map, which determines where its trajectories end. Replay thus may be how that structure is kept current as the task changes.

## Methods

### Default representation

#### Linear RL

Our model builds on the framework introduced in the linear RL model [18]. The linear RL model provides a closed-form approximation to the optimal value function for finite-horizon Markov decision problems, such as maze navigation towards a specific goal. At its core, this model optimizes behavior by maximizing a gain function that balances rewards against control costs [49]. The gain function *g* for a state *s* is defined as:

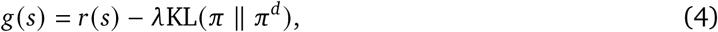

where *r*(*s*) represents the reward at state *s*, and *λ* > 0 is a control cost parameter. The second term measures the divergence between two policies: the agent’s decision policy *π* and a default policy *π^d^* under which the predictive map is built. This divergence is quantified using the Kullback-Leibler (KL) divergence, which equals zero only when the policies match exactly (*π* = *π^d^*) and is positive otherwise. Throughout our simulations, we use a uniform default policy where all transitions to neighboring states are equally likely. A uniform random default policy introduces stochasticity into decision-making without biasing the decision policy toward any specific actions. It can be shown that the optimal value function *V*^∗^(*s*) for this problem has an analytical solution, which can be viewed as an approximation to the value function of the original RL problem without control costs. To derive this solution, we first define the one-step state transition matrix **T**. Each element *T_i j_* represents the probability of transitioning from state *i* to state *j* under the default policy.

Next, we partition the state space into terminal states (containing goals) and nonterminal states. Let **t** = **T***_G_* exp(**r***_g_*/*λ*) be a vector determined by transitions from nonterminal to terminal states **T***_G_* and their associated rewards **r***_G_*. For a vector **v**^∗^ containing optimal values across all nonterminal states, we have:

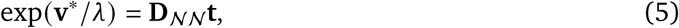

where **D**_NN_ is the submatrix of the DR matrix **D** corresponding to nonterminal states. Recall that the DR matrix is defined as:

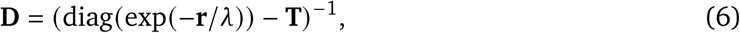

where **r** is the reward vector across all states (assuming non-zero rewards at terminal states). Thus the DR, **D**, is a matrix where the (*i*, *j*) element indicates the expected occupancy of state *j* from state *i* discounted by exponential state rewards (which are negative) when following the default policy.

##### Woodbury update

The second critical component introduced in conjunction with the linear RL framework was the Woodbury update, a low-rank and easy to compute update to the predictive map when there is a change in the environment [18, 19]. This contrasts with both classic model-based approaches and the SR, which typically require recalculating the entire map even for minor environmental changes.

Formally, when an environmental change modifies matrix **L** = diag(exp(−**r**/*λ*)) − **T** to a new matrix **L**_new_ (due to alterations in either **T**, **r**, or both), we can apply the Woodbury matrix inversion lemma. Given **D** = **L**^−1^, this lemma allows us to compute the new matrix **D**_new_ = **L**^−1^ through a computationally efficient low-rank update of **D**.

Now, let us introduce two sparse matrices **R** and **C** of size *J* × *S* and *S* × *J* respectively, where *S* is the total number of states and *J* is the number of states for which **L** has changed. Let **j** = ( *j*_1_, *j*_2_, …, *j_J_*) be a vector of length *J* containing the indices of all those states, where *j_i_* denotes the *i*-th element of the vector. For each changed state, matrix **R** contains rows representing the difference between the corresponding rows in **L**_new_ and **L**, such that **R** = **L**_new_(**j**, :) − **L**(**j**, :). Matrix **C** acts as a selection matrix: for each changed state *j_i_*, it contains a column with a value of 1 in row *j_i_* and zeros elsewhere. Using these matrices **C** and **R**, we can efficiently express the change in **L** as:

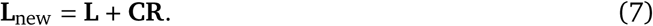

Using this formulation, we can apply the Woodbury matrix inversion lemma [50, 51] to express

**D**_new_ (the inverse of **L**_new_) as a low-rank update of **D**:

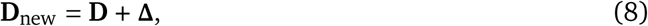

where the change matrix **A** is given by

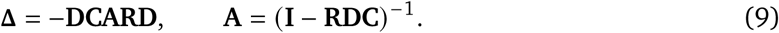

Importantly, while this formulation still requires matrix inversion for replanning, it operates on a much smaller matrix: **A** has dimensions *J* × *J*, where *J* is the number of changed states and is typically much smaller than the total number of states *S*. Equation (8) gives the exact repair, applied to every row of **D** at once. We treat this as the target that replay approximates, not as the operation replay carries out; the state-wise implementation (equation (3) is described in more detail next.

### The map-repair model of replay

We propose that replay is the process by which the map is repaired after a goal switch, and that the two corrections a goal switch requires correspond to prospective and retrospective replay. Each time the map must be repaired to plan from one goal to the next, our model applies two separate updates, in two steps. Let **D** be the map in place before either has been applied. Both updates are computed from **D** using equation (9): **Δ***_P_* makes the new destination terminal, and **Δ***_R_* makes the previous destination nonterminal. Each is a low-rank change matrix applied to the predictive map as in equation (8). Neither is evaluated on a map to which the other has already been applied, so each treats the change it concerns as the only change. This is exact when the two changes are far enough apart that their effects on the map do not interact. When they are close, their effects no longer decompose into a simple sum, but the pair remains a principled first-order approximation to the exact correction, as for the composition of multiple objects [19].

#### Which repair is carried out first

We identify prospective replay with making the new goal terminal, **Δ***_P_*, and retrospective replay with making the previous goal nonterminal, **Δ***_R_*. Switching the goal from *g*_old_ to *g*_new_ changes two rows of **L**: making *g*_new_ terminal sets row *g*_new_ of **T** to 1 on the diagonal and 0 elsewhere, so that the state transitions only to itself and no path continues beyond it, and making *g*_old_ nonterminal restores its default row, allowing transitions out of it again. Each is a *J* = 1 change, so **A** of equation (9) is a scalar and each change matrix is rank one. Its single nonzero eigenvalue is *λ* = −**A RD**^2^ **C**, which gives a precise definition of the magnitude of a repair: how much of the map it changes.

The two repairs are carried out in the order of their magnitudes, with the larger first. We select between them by their proportional probability, that is, the probability of carrying out the prospective repair first is |*λ_P_* | relative to the sum of the two magnitudes, where *λ_P_* and *λ_R_* are the eigenvalues of **Δ***_P_* and **Δ***_R_* respectively.

#### How each repair is applied

Recall that neither repair is added to **D** in a single step. Instead, a replay event starting at the animal’s current state *s*_0_ traces a trajectory *s*_0_, *s*_1_, … , *s_K_* through the state space and applies equation (3) at each visited state. The states whose rows are corrected are therefore not states the animal occupies, which is what makes the repair an offline operation: it corrects the map where the animal is not. The next state is chosen deterministically from the update matrix itself: from *s_k_* the replay moves to the adjacent, unvisited state *s*^′^ that maximizes |**Δ**(*s_k_*, *s*^′^) |, terminating when it reaches the state whose terminal status the update concerns. Rows of states not visited by the replay are left unchanged. The update is therefore exact along the replayed path and absent elsewhere, so which states are replayed, and in what order, determines which parts of the map are corrected. Because the walk ends at the state being changed, the replayed trajectory is a path between the animal’s current location and the goal that update concerns.

What we are left with are two separate updates to the predictive map along the path of each replay’s respective trajectory, **Δ***_P_* making the new destination terminal and **Δ***_R_* making the previous destination nonterminal. The two repairs are applied one after another to the predictive map **D**, in the order of their magnitudes. If the repair associated with **Δ***_P_* is the larger, we first apply the terminal repair associated with *g*_new_ and then the nonterminal repair associated with *g*_old_:

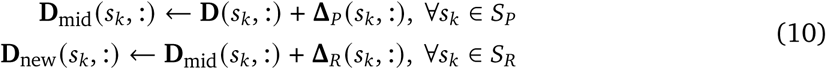

Here **D** is the map in place before either repair, **D**_mid_ is the map after the first repair has been applied, and **D**_new_ is the map after both. *S_P_* and *S_R_* are the states along the prospective and retrospective repairs respectively.

#### Direction within a repair

In many replay experiments, especially those in track environments (linear track, T-maze, W-maze), place cells do not behave as they do in an open field: they become aligned with a specific direction of travel along the track. Replay in these studies is therefore reported both as a type (prospective or retrospective) and as a direction.

To model this, recall that a change matrix **Δ** is the same size as the map, so it assigns a value to every pair of states. Positioning ourselves at a chosen state *s* within the repair, the entries of **Δ**(*s*, :) give the size of the correction along each path leading away from *s*, and the relative size of these entries gives the probability of the repair proceeding in each direction. For two candidate states *s*_1_ and *s*_2_, the probability that the repair proceeds towards *s*_1_ is |**Δ**(*s*, *s*_1_) | relative to the sum of the two. Directions that require a larger correction are therefore replayed more often. Because the repair is exact, the model specifies the relative weight of each path rather than a number of replay events, and we report these relative weights throughout.

In sum, when we simulate studies that probe the direction of replay, we position ourselves inside the repair and report the probability of each direction as the proportion of replay in that direction. When we are focusing on replay as a whole, prospective versus retrospective, we report the probability derived from the eigenvalues of the two repairs, or in the case of Ambrose et al. [9] the percent difference between them. The corrections themselves are exact, so treating their relative sizes as probabilities is a simplifying assumption: the proportion of replay in a given direction reflects both which repair is selected and where that repair proceeds once selected.

### Simulation details

#### Environments and state spaces

Throughout the study, we consider the environments to be tabular of shape *n_r_* × *n_c_* where *n_r_* is the number of rows and *n_c_* is the number of columns. The agent can move in each of the cardinal directions. When “selecting” the replay path from the change matrix, **Δ**, we assume that each state is chosen deterministically by the largest value. Thus, if we are “inside” of the replay update and the update is currently at state *s*, the next state is chosen as argmax*_s_*′ **Δ**(*s*, *s*^′^). Across all of the simulations, unless otherwise stated, we fix the reward at all nonterminal (*r*_N_) to be −0.1 and set *λ* to 1.

#### Efficiency of the state-wise update

For the proof of concept in showing that our method can replan optimally (Supp. Figure 1) on-par with the entire matrix version of the repair, we set the reward at the terminal state to be 10. We initially set the terminal reward state to be in the bottom-right of the maze (r_1_) and use the closed-form solution from equation (1) to get the optimal DR with respect to that terminal state, which gives us **D**. We then move the terminal state to either the bottom-left (s_1_) or the top-right (s_2_) of the maze and get the associated change matrix. We now have two updates: one to make the previous goal location (r_1_) nonterminal and another to make the new goal location (s_1_ or s_2_) terminal. We update the representation using the aforementioned replay state selection method to get **D**_new_. We then use the updated representation to get a value function (equation (5)) and use that to plan towards the goal. We average across 50 seeds.

#### Ordering of repairs and its dependence on path geometry

For our modeling of Mallory et al. [6], we generate 12 5 × 5 mazes, identical to ones that were used in the original study. Following their design we place 9 reward wells in the middle 3 × 3 section of the maze. For each maze we simulate 50 sequences, with random seeds fixed across the mazes. Each sequence begins with a randomly sampled home well, followed by 3 away wells, each drawn from the wells other than home and different from the one before it, and the agent alternates between home and the away wells as in the original task. We sample 3 away wells which results in 6 trials per sequence. With our home state and 3 away states we then run through a series of “home” and “away” trials, following the task design of the original study. At every goal switch, the prospective repair (making the next goal terminal) and the retrospective repair (making the current goal nonterminal) are computed from the current map and applied along their replay paths (equation (10)). The map is carried forward from one switch to the next, so it reflects the sequence of repairs that preceded it.

Because the repair magnitudes are a deterministic function of the map and the two goals, the map can be probed directly: at any point in the sequence, we can evaluate the repairs for a given choice of next goal without carrying them out. We use this to compare converging and diverging configurations on the same map, so that the comparison is not confounded by differences in the history of repairs. At each switch to a new away well, we evaluate the repairs for that well, which always differ from the well the agent just came from (diverging trials), and, on the same map, for the well the agent has just come from as the next goal, in which case the past and future paths coincide (converging trials). Only the repairs for the actual next well are applied, so the sequence of maps is the same in both cases, and the two configurations are defined by the goals rather than by a threshold on path angle. For comparing between prospective and retrospective directly across both trial types, we average the probabilities of the prospective repair being carried out first across the two conditions, different sequences, and different mazes. For our analysis of retrospective replay on converging and diverging trials, we compute, for each trial type, the optimal distance from the well the agent has just come from to its current location, the number of steps on the shortest path between them given the maze walls, and plot the probability of retrospective replay at each distance from 1 to 5 steps.

#### Arbitrating between prospective and retrospective updates

The modeling of Diba and Buzsáki [2], and Ambrose et al. [9] both involve arbitrating between prospective and retrospective updates (Supplementary Figures 2 and 4). Again, in our model we assume that making the future location terminal is a prospective update while making the previously terminal location nonterminal is a retrospective update. Both are invoked together to accommodate efficient goal-directed planning from one goal to the next. We follow a similar approach as described above but now applied to a linear track environment. For Diba and Buzsáki [2] we make one end of the track terminal and compute the probability of prospective vs. retrospective replay as described above. For Ambrose et al. [9], we are focused on the event when an agent reaches one end of the track and must make the currently reached state nonterminal and the other end terminal. Along with the terminal and nonterminal updates, we also modify the reward of the state that the animal reached (as they do in the paper) with the 0x condition being no reward, 1x being a reward of 1, and 4x being a reward of 4. We use the eigenvalues of the Woodbury updates to model the relative weights of each replay update and take the percent difference between the eigenvalues of each 4x and 1x condition as well as between the 0x and 1x conditions to get the values we plot.

#### Directionality within an update

The modeling of Shin et al. [8], Carey et al. [13], Gillespie et al. [12], and Ólafsdóttir et al. [10] all involve the directionality within a repair specifically. These studies do not focus so much on arbitrating between prospective and retrospective replay as much as they are focused on specifically *where* the replay is going. Our model naturally extends to that and we compute the probability of each direction as described above.

For Shin et al. [8], we construct a w-maze (see Figure 3**a** for a diagram). We categorize inbound and outbound paths in the same manner as the authors: inbound paths are the previous path the rat/agent took to get to their current location and outbound paths are the future paths the rat/agent will take to get to the endpoint of the next arm (well). For our simulation we start with the base DR of the environment and initially set the agent to be at either a side well or the center well, depending on whether we are simulating the “center well” or “side well” condition respectively. We then perform a repair update to make the center well (if in a side well) or one of the side wells (if in the center well) terminal and apply that update to get a new predictive map. The agent has now moved into either the center well or one of the side wells. Again we make either the side well or center well terminal, but this time we also make the well that the agent is currently at nonterminal, so that we have two replay conditions that we can categorize as prospective and retrospective respectively. Because we are interested in the directionality within the replays specifically, we position ourselves “within” the replay at the bottom-center of the maze. This allows us to compute the probability of the replay going towards the inbound path or the outbound path, as described above.

For Gillespie et al. [12], the agent was placed at the start of an 8-arm maze (see Figure 4**a** for a diagram). We make two assumptions. The first is that once an arm has been found to be rewarding, that state must be made terminal in order for the “repeat” phase to take effect. The rats, in theory, learn that once a reward is found it will not move, so it makes sense for this to be stitched into the cognitive map for long-term use. Secondly, we assume that when the rat has found the arm to be unrewarding, thereby prompting the “search” phase of the task, the currently terminal arm must be made nonterminal as it is no longer rewarding. We model the task by computing the DR initially assuming that some arm is rewarding and thus terminal, so that our replay update is to make that arm nonterminal. We plot the probability of the replay expanding from the start of the maze into each separate arm.

For the T-maze tasks of Carey et al. [13] and Ólafsdóttir et al. [10] we use the same environment but with slightly different task structures. In the case of Ólafsdóttir et al. [10], we model each arm as being non-terminal, while in the case of Carey et al. [13], we model each arm as being terminal. This choice is not arbitrary and points to a deeper question about task schema and how an animal might encode a representation of the environment for planning. In the case of Ólafsdóttir et al. [10], the animals are not allowed to run beyond the junction of the t-arm, making either arm inaccessible and thus non-terminal. It is only at test that the barrier is removed and the rewarding arm is marked with a cue for the rat. We argue that at this point it would make sense for the rat to make the cued arm terminal as it wants to plan towards it and thus we see the classic “goal-directed” replay. Whereas in the case of Carey et al. [13] both arms are accessible and each contains a unique reward (either food or water). The rat learns which arm contains what and they are allowed to come and go, getting as much food and water as they please. We argue in this setting each arm is its own unique terminal state. Only at test, when the rat is restricted access to a resource and placed back into the maze, does it make sense for the rat to make the arm it does not want to travel to nonterminal. And thus we see “paradoxical” non goal-directed replay. For both situations we take the base map with both arms nonterminal or terminal and get the repair to make one arm terminal or one arm nonterminal. We then position ourselves at the junction in the T-maze and compute the probability of replay going towards each arm, as described above.

### Analysis of hippocampal recordings

#### Dataset

We analyzed hippocampal recordings previously reported in Mallory et al. [6]. Full surgical, recording, spike-sorting, and behavioral procedures are described there. Briefly, the spiking activity of large ensembles of place cells was recorded from the bilateral dorsal hippocampus of four rats navigating a 90 × 90 cm open field arena. Rats were rewarded on alternating trials at a consistent, learnable location (“home” trials) or at an unpredictable location that varied from trial to trial (“away” trials), with a mean±s.e.m. of 88 ± 4 trials per session across all sessions. A mean±s.e.m. of 223 ± 10 cells were co-recorded per session. In the majority of sessions (*n* = 38), the arena contained transparent barriers, which the animal was required to circumnavigate. The present analyses used 44 sessions from 4 rats, comprising 1908 away trials that met the inclusion criteria below.

#### Replay detection and classification

Brief bursts of synchronized pyramidal cell activity (spike density events; SDEs) were considered candidate replay events. To identify SDEs, total spike activity across all putative pyramidal cells was computed in 1-ms bins throughout the session and z-scored across periods of immobility (movement speed < 5 cm/s). Times at which the z-scored spike activity exceeded 3 standard deviations were identified as SDE peaks, with event onset and offset defined as the times at which the z-scored activity returned to zero on either side of the peak. Within each SDE, position was decoded from ensemble spiking using a memoryless Bayesian decoder with a uniform spatial prior, in 20-ms bins advancing 5-ms steps. Candidate events lasting > 50 ms, recruiting > 10% of pyramidal cells, and spanning > 20 cm from start to finish were classified as replay events.

For each replay, we computed the angular displacement between the replayed trajectory and the animal’s immediate past path (RP) and immediate future path (RF), each defined over the 100 cm of trajectory adjoining the stopping location. Angular displacement was measured by centering a series of concentric rings on the animal’s stopping location, beginning at a radius of 14 cm and increasing in 2 cm increments. At each radius we identified the intersections of the ring with the replay trajectory and with the relevant behavioral path (past or future) and calculated the absolute angular separation between these intersection points relative to the animal’s location. Angular displacement was then averaged across radii at which both trajectories intersected the ring to obtain a single RP or RF value for each replay. Replays with RP < 20^◦^ and RF > 20^◦^ were classified as retrospective, whereas replays with RF < 20^◦^ and RP > 20^◦^ were classified as prospective, following Mallory et al. [6]. All analyses reported here were restricted to retrospective replay events.

#### Trial variables and inclusion criteria

Analyses were restricted to away trials, on which the rewarded well was drawn at random and so the geometry relating the previous, current and next wells varied from trial to trial. We restricted our analysis to away trials because only one of these trials can the animal know where the next goal location is (Home). For each away stopping period, we computed: the past–future path angle (PF), the angular displacement between the animal’s immediate past and immediate future paths, computed exactly as in Mallory et al. [6] (their Figure 3B); the optimal path length from the previous well to the current well, defined as the length of the shortest traversable path between them given the barrier configuration in that session. Trials were classified as having converging past and future paths if PF < 60^◦^ and diverging paths if PF > 120^◦^; these thresholds are those used in Mallory et al. [6] and were not adjusted here. Of the 1908 away trials entering the analysis, 925 were classified as converging and 339 as diverging, with the remainder falling between the two thresholds and excluded from the split analyses.

#### Retrospective replay rate

For each stopping period, retrospective replay event rate was defined as the number of retrospective replays divided by the stop duration, expressed in Hz.

#### Binned replay rates (Figure 5c,d)

Within each path-geometry class (converging or diverging), optimal distances were divided into five equally spaced bins spanning the observed range (20–160 cm). For each bin we computed the mean retrospective replay rate across the stopping periods falling in that bin; error bars denote s.e.m. across stopping periods.

#### Poisson regression

To test the model’s prediction statistically, we modelled the number of retrospective replay events emitted during each stopping period as a Poisson random variable with log(stopping period duration) as an offset, so that coefficients are interpretable as effects on replay rate rather than count. Models were fit by maximum likelihood (fitglm, MATLAB R2025b). We first asked whether the effect of distance depends on path geometry by fitting, across all away trials, the model

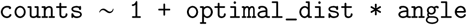

with angle entered as a continuous variable in radians (*n* = 1908). We then fit the reduced model counts ∼ 1 + optimal_dist separately within the converging (*n* = 925) and diverging (*n* = 339) subsets. To confirm that the results were not an artifact of pooling stopping periods across animals and recording sessions, each model was refit as a generalized linear mixed-effects model (fitglme) with a random intercept for rat, and again with a random intercept for unique recording session.

#### Non-parametric confirmation

As a complement to the parametric models, and to verify that the results did not depend on the assumed linear form of the distance effect, we performed two additional tests within each geometry class. First, we split trials at the 33rd and 66th percentiles of the optimal-distance distribution (computed across all trials) and compared mean retrospective replay rates between the closest and furthest terciles using a two-sided Wilcoxon rank-sum test. Second, we computed the Pearson correlation between optimal distance and mean retrospective replay rate across all trials in the class.

## Data availability

All new data in this study were obtained through simulation. This study also re-analyzed existing data from Mallory et al. [6], which are available from C.S.M. upon request.

## Code availability

Analyses were performed using custom code developed in Python and MATLAB. Code for the model, simulation, and analysis is available at https://github.com/arminbazza/Replay.

## Notes

### Competing Interest Statement

The authors have declared no competing interest.

### Summary of Updates

We added middle initials to two author names and slightly revised the manuscript.

