## Supplementary material for "Hippocampal replay repairs the cognitive map for flexible planning"

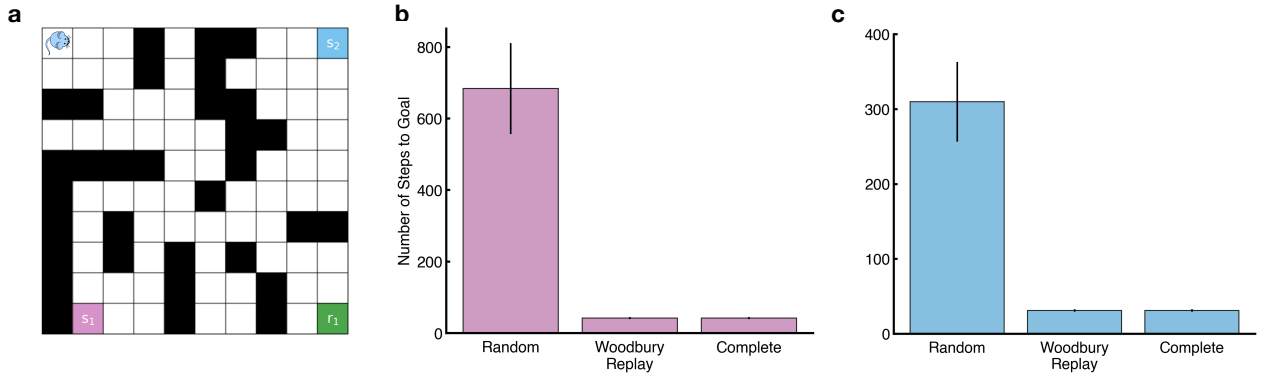

Figure 1: **Stepwise repair preserves planning**

We validate the utility of stepwise repair for planning. We test whether stepwise repair updates the predictive map well enough to plan optimally to a new goal elsewhere in the environment.

(a) Maze we tested the model in. Initially the model learns a representation of the environment (maze) assuming that $r_1$ is the only rewarding terminal state. At test, we place the agent in state $s_1$ and then update the representation to make either state $s_2$ or $s_3$ the new rewarding terminal state and set $s_1$ to now be nonterminal. We then test the agent’s path length from $s_1$ to both $s_2$ and $s_3$. We compare the agent that updates its representation stepwise with the agent that completely reconstructs a new representation (Complete). We also include a completely random agent as a baseline.

(b) Replanning from $s_1$ to $s_2$. The agent updating its representation stepwise is nearly as optimal as the Complete agent.

(c) Replanning from $s_1$ to $s_3$. Again, the agent updating its representation stepwise is nearly as optimal as the Complete agent..

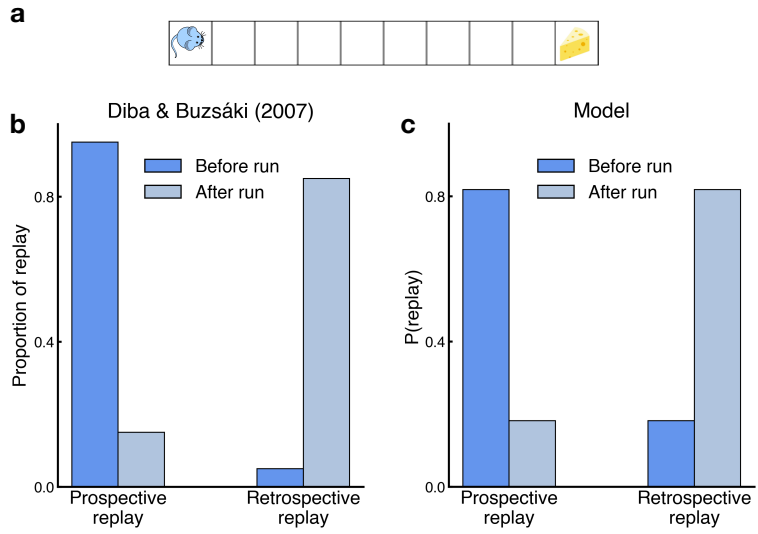

Figure 2: **Forward and reverse replays on a linear track**

(a) Linear track where the rat runs from end to end, receiving a reward at each end of the track.

(b) Data from Diba and Buzsáki (2007) showing that the majority of forward sequences occurred at the start end of the track before running, while the majority of reverse sequences occurred at the other end following the run. Data is plotted as proportion of replay events instead of raw number of events to match current approaches with plotting replays.

(c) Forward replay is more likely to occur prior to the run, reverse replay is more likely to occur after the run with its probability being modulated by the probability of forward replay initially being selected.

Image in (b) adapted from (Diba and Buzsáki, 2007), Nature Publishing Group.

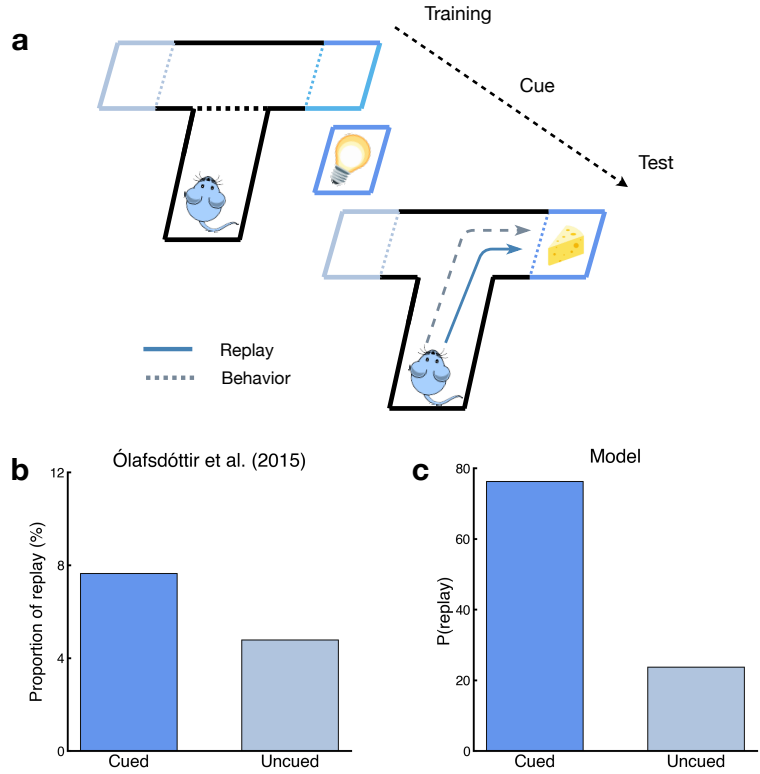

**Figure 3: Goal directed replay in a T-maze**

**(a)** T-maze environment. The rats were initially placed in the bottom of the maze and could only run up towards the junction where the maze split into the left and right arm. One of the two arms was then cued, indicating to the rat which arm contained a reward. The barrier was removed and the rat was allowed to run towards the arm. Replays were recorded prior to the commencement of the run.

**(b)** Data from Ólafsdóttir et al. (2015) showing the proportion of replay events going towards the cued and uncued arms respectively. The majority of events go towards the cued arm where the goal is.

**(c)** The majority of replays from the model goes towards the cued arm which contains the goal as opposed to the uncued arm.

Image in **(b)** adapted from (Ólafsdóttir et al., 2015), Creative Commons License CC BY 4.0.

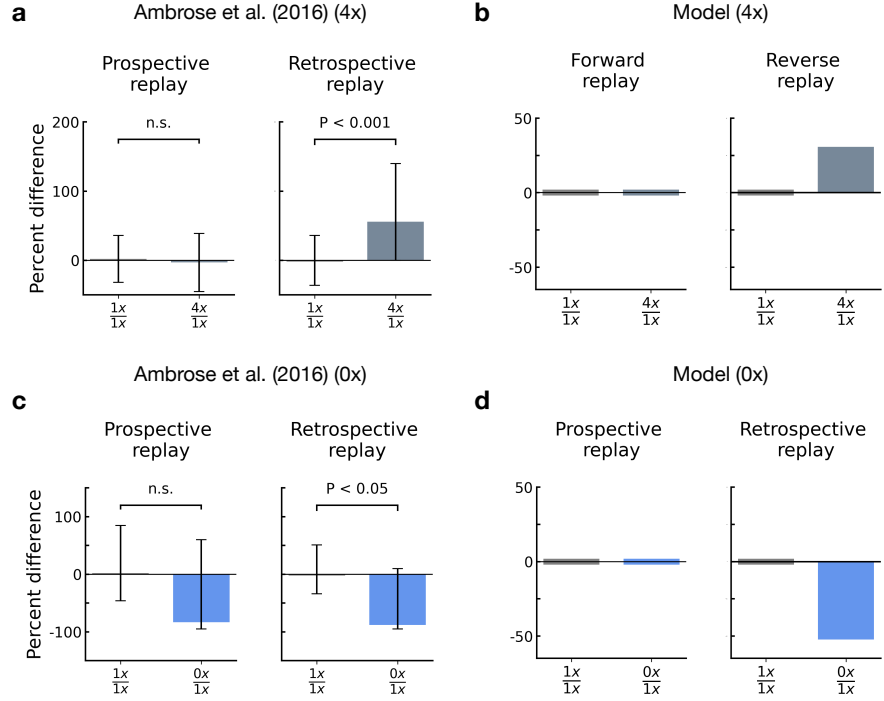

Figure 4: **Impact of reward modulation on forward and reverse replay**

(a,b) Effect of increasing the reward to 4x: (a) Replotted data from Ambrose et al. (2016) showing percent difference in replay rate from unchanged to increased reward at one end of the track. (b) Simulation of our model on the task showing no difference in the eigenvalue of forward replay (left) and an increase in the eigenvalue of reverse replay in the 4x condition (right, grey).

(c,d) Effect of decreasing the reward to 0x: (c) Replotted data from Ambrose et al. (2016) showing percent difference in replay rate from unchanged to decreased reward at one end of the track. (d) Simulation of our model on the task showing no difference in the eigenvalue of forward replay (left) and a decrease in the eigenvalue of reverse replay in the 0x condition (right, blue).

Image in (a,c) adapted from (Ambrose et al., 2016), Elsevier.
